# Development of transcriptomics-based eukaryotes growth rate indices

**DOI:** 10.1101/2021.10.08.463626

**Authors:** Wye-Lup Kong, Ryuji J. Machida

## Abstract

1. Growth rate estimation is important to understand the flow of energy and nutrient elements in an ecosystem, but it has remained challenging, especially on microscopic organisms.
2. In this study, we propose four growth rate indices that use mRNA abundance ratios between nuclear and mitochondrial genes: (1) total nuclear and mitochondrial mRNA ratio (Nuc:Mito-TmRNA), (2) nuclear and mitochondrial ribosomal protein mRNA ratio (Nuc:Mito-RPmRNA), (3) gene ontology (GO) terms and total mitochondrial mRNA ratios and (4) nuclear and mitochondrial specific gene mRNA ratio. We examine these proposed ratios using RNA-Seq datasets of *Daphnia magna* and *Saccharomyces cerevisiae*.
3. The results showed that both Nuc:Mito-TmRNA and Nuc:Mito-RPmRNA ratio indices showed significant correlations with the growth rate for both species. A large number of GO terms mRNA ratios showed significant correlations with the growth rate of *S. cerevisiae*. Lastly, we identified mRNA ratios of several specific nuclear and mitochondrial gene pairs that showed significant correlations.
4. We foresee future implications for the proposed mRNA ratios used in metatranscriptome analyses to estimate the growth rate of communities and species.

## 1 | INTRODUCTION

Organisms’ growth rates are important traits that can be influenced by environmental conditions of its habitat, such as temperature and availability of nutrients (Logan et al., 1976; Urabe et al., 1997; McCarthy et al., 1998). By estimating the growth rate of the organisms in an ecosystem, we can determine the efficiency of energy and nutrient element flow to higher trophic levels in a food web system (Cloern et al., 1995; Granados et al., 2017). Growth rate varies between species and population, and changes according to the spatial and temporal variation in environmental conditions. Hence, an effective growth rate estimation method is essential to understand ecosystems.

Growth rate estimation remains challenging, especially for microscopic organisms, because individual tracing and direct measurement methods are not feasible. Hence, approaches such as biochemical indices are favoured in growth rate estimation. Over the decades, many nucleic acid-based indices have been introduced to estimate the growth rate of an animal population, many of which use RNA content as the proxy for growth rate (Yebra et al., 2017). There are three main reasons the nucleic acid indices have gained popularity in growth rate estimation. First, the approach does not require the incubation of animals, where the samples can be collected from the field and the nucleic acid quantified to estimate the growth rate. Second, the method is applicable across different taxonomic groups due to the conservative feature that mRNA is being used as the blueprint of protein production (Crick, 1970). Third, molecular techniques are easy to access because of the availability of commercial kits. These kits can reduce the time required for training and improve the quality of results.

Initially, RNA concentration was considered as the proxy for animal growth rate (Sutcliffe, 1965). After two decades, Ota and Landry (1984) followed up with animals’ conservative feature that uses the RNA:DNA ratio indices with DNA concentration as the denominator. The RNA:DNA ratio gained popularity and was widely incorporated in many studies across different taxonomic groups due to its simple and rapid procedure (Wagner et al., 1998). However, Ota and Landry (1984) also pointed out the limitation of the ratio used in field-collected samples. One of the reasons might be the size of the genome, which can vary drastically among eukaryote species (Gregory, 2007). Another novel nucleic acid index, the nuclear to mitochondrial ribosomal ratio (Nuc:Mito ribosomal ratio), uses quantities of cytosolic and mitochondrial ribosomal RNA (rRNA) as numerator and denominator, respectively, for the ratio (Kong et al., 2019). The significant correlation between the ribosomal ratio and growth rate is due to the allocation of assimilated nutrient elements to the mitochondrial and nuclear ribosome, responsible for energy production and growth-related genes, respectively. The usefulness of the ratio suggested that the overall gene expression levels between nuclear and mitochondrial genomes might reflect the growth rate of eukaryotic organisms.

Over the past decades, high-throughput transcriptomics technologies have gradually been incorporated into ecology studies (Alvarez et al., 2015; Caron et al., 2017; Brauer et al., 2017). Through transcriptomics analyses, researchers can capture the gene expression responses towards the environmental changes and compare different locations with various conditions. In this study, we investigated the potential of four mRNA ratio indices: (1) total nuclear and mitochondrial mRNA ratio, (2) nuclear and mitochondrial ribosomal protein mRNA ratio, (3) gene ontology (GO) terms and total mitochondrial mRNA ratio and (4) nuclear and mitochondrial specific gene mRNA ratio. Transcriptome datasets of *Daphnia magna* and *Saccharomyces cerevisiae* were used for the study. The results demonstrated that the proposed mRNA ratios are useful growth rate indices, which might be used for individuals and communities collected in the field.

## 2 | MATERIALS AND METHOD

This study included two different transcriptome datasets to evaluate the proposed nuclear and mitochondrial mRNA ratios. (1) The first transcriptome dataset is the freshwater water-flea (*Daphnia magna*) cultured at different temperatures and food concentrations. (2) The second dataset is the *S. cerevisiae* transcriptome dataset, cultured with different phosphorus-level treatments, which was retrieved from the National Center for Biotechnology Information (NCBI) (accession number: PRJNA395936, Gurvich et al., 2017).

### 2.1 | *Daphnia magna* acquisition and preparation

*Daphnia magna* samples were obtained from the Freshwater Bioresources Center, National Chiayi University, Chiayi, Taiwan. The samples used in this study were cultured in laboratory incubators under 16:8 hours light and dark cycle. Each sample was kept individually in a 50 ml tube filled with dechlorinated and filtered tap water. Samples were fed with laboratory-grown green alga *Chlorella vulgaris* as a food source and fed daily by replacing the culturing water with the green alga. Different levels of temperature (High: 25, Medium: 20 and Low: 15 °C) and food concentration (High: 5 × 10^5^, Medium: 5 × 10^4^ and Low: 5 × 10^3^ cells/ml) were used to manipulate the growth of *D. magna*. Each treatment group consisted of five replicates. The body length was recorded twice, newly hatched (start of the incubation) and immediately before the development of the embryo (ovary becomes darker and swollen: end of the incubation). Dry weights of the individuals were estimated based on a length–weight regression formula (Kawabata & Urabe, 1998; Weight = e^3.05 + 2.16ln(Length)^). The somatic growth rate of *D. magna* was calculated based on the formula provided by Lampert and Trubetskova (1996):

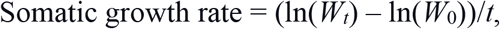

where *W_t_* is the weight (μg) of the sample collected for RNA extraction, *W*_0_ is the sample’s weight on the first day of culture and *t* is the time culture duration (day).

### 2.2 | Ribonucleic acid extraction

Total RNA was extracted using the RNeasy Plus Micro kit (Product ID: 74034, QIAGEN, Hilden, Germany) and followed the standard protocol provided by the manufacturer.

### 2.3 | High-throughput transcriptome

#### 2.3.1 | Transcriptome library preparation and sequencing of *Daphnia magna* mRNA

Twenty ng of the total RNA were used for each library preparation. The mRNA was isolated from the total RNA using the NEBNext Poly(A) mRNA magnetic isolation module (Product ID: E7490, New England Biolabs, MA, USA), following the standard protocol provided by the manufacturer. Transcriptome library preparation procedures were performed following the standard protocol of NEBNext Ultra II RNA Library Prep Kit for Illumina (Product ID: E7770, New England Biolabs) with some modifications. The enrichment step for the library preparation was extended from 20 to 25 cycles to ensure an adequate amount of DNA for the sequencing. The Illumina MiSeq (Illumina, CA, USA) with 150 cycles single read was used for the sequencing. The NGS Core Laboratory in the Biodiversity Research Center, Academia Sinica, Taiwan conducted all sequencing procedures.

#### 2.3.2 | Sequence data analysis

Standard low-quality reads filtering and quality checks were done using the program cutadapt (Martin, 2011) and fastqc (Andrews, 2010), respectively, with a minimum sequence length of 50 bp. Filtered high-quality sequences were mapped to the reference genome using the program Tophat (Trapnell et al., 2009). Reference genome (fasta format) and general feature file (gff format) for *D. magna* were retrieved from wFleaBase (wfleabase.org; Colbourne et al., 2005). A maximum of five base pair mismatches was allowed during the mapping process. HTseq (htseq-count; Anders et al., 2015) was used to acquire sequence read counts for both nuclear and mitochondrial protein-coding genes. During the read counting process, only sequences mapped to protein-coding genes were included. Ratio indices, correlation coefficient (function: cor.test) and adjusted coefficient of determination (function: lm) were calculated using the R platform (R Core Team, 2020). All the commands used in the process are provided in the supporting information (Supp. Info 1).

#### 2.3.3 | Ratio indices calculations

There are four mRNA ratio indices proposed in this study.

1. Total nuclear and mitochondrial mRNA ratio (Nuc:Mito-TmRNA)
2. Nuclear and mitochondrial ribosomal protein mRNA ratio (Nuc:Mito-RPmRNA)
3. GO terms and the total mitochondrial mRNA ratio
4. Nuclear and mitochondrial specific gene mRNA ratio

The first ratio, Nuc:Mito-TmRNA, uses the abundances of mRNA reads mapped to the reference of the nuclear genome as the numerator and the reads mapped to the mitochondrial genome as the denominator. The second ratio, Nuc:Mito-RPmRNA, uses the number of mRNA reads mapped to nuclear and mitochondrial ribosomal protein-coding genes. The third ratio is the GO term and mitochondrial mRNA ratio, which uses the reads numbers mapped on the lowest child GO terms and mitochondrial mRNAs. GO information from the UniProt database (uniprot.org, 2021) for the *D. magna* dataset was utilized for the annotation. Manual annotations were performed by pairing the protein name of genes in our dataset with the protein list in databases. Genes were grouped based on the annotated GO terms and each GO read abundance was calculated. In the ratio, total mitochondrial mRNA read abundance was used as the denominator. The last mRNA ratio uses read abundances of a single specific nuclear and a mitochondrial gene read abundance. Every nuclear and mitochondrial gene combination available in the transcriptome dataset was calculated in this study. All the mRNA ratios were temperature corrected using the van’t Hoff–Arrhenius relation because most of the biological processes rates are influenced exponentially by temperature (Brown et al., 2004),

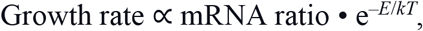

where *E* is the activation energy (0.63 eV), k is Boltzmann’s constant (8.62 ×10-5 eV· K-1) and T is the temperature in kelvins. The value of activation energy was derived from the temperature-correction of metabolic rate (Brown et at., 2004). All the calculations were done on the R platform. The R scripts for the calculation procedure are uploaded into Zenado Repository (https://doi.org/10.5281/zenodo.5506660).

### 2.4 | Real-time quantitative PCR

To validate the nuclear and mitochondrial specific gene mRNA ratio, we performed the absolute quantification approach of real-time qPCR on the top 10 nuclear and mitochondrial gene pairs that have the highest correlation coefficients with growth rate.

#### 2.4.1 | RNA extraction and reverse transcription for real-time quantitative PCR

RNA extraction was performed using PureLink RNA Minikit (Product ID: 12183020, Invitrogen, CA, USA). Reverse transcription was performed using Maxima Reverse Transcriptase enzyme (Product ID: EP0741, Thermo Fisher Scientific, CA, USA) together with random hexamer primers (Product ID: N8080127, Invitrogen).

#### 2.4.2 | Primer design

Two types of primers were designed for the real-time qPCR experiment. The first primers, known as the precursor primers, amplified the target genes’ longer sequence product (ca. 1,000 bp). The purpose of these amplified products is to construct standard curves that can be used in absolute qPCR. The second set of primers is designed based on the precursor primers’ product sequences. These primer sets are used in the real-time qPCR. All the primers used in this study were designed using the program Primer3 (Untergasser et al., 2012) and listed in the supporting information (Supp. Table 1).

#### 2.4.3 | Real-time quantitative PCR

Absolute real-time qPCR was conducted using SYBR-green PowerUp master mix (Product ID: A25776, Applied Biosystems, MA, USA) following the standard protocol. QuantStudio 3 Real-Time PCR system (Applied Biosystems) was used in the PCR. All cDNA template concentrations were standardized to 1 ng/μl, and primers’ concentrations were diluted to 3 mM. Each qPCR plate consisted of the test cDNA samples, a 5-point standard curve (1 × 10^9^ to 1 × 10^5^ copy numbers), and a no-template control for each primer set. Every sample was loaded into three wells as triplicates during the qPCR process. The standard curve of each qPCR plate was used as a quality filter with the criteria of primer efficiency ranging from 110% to 90% and *R*^2^ value higher than 0.98.

#### 2.4.4 | Ratio indices calculations

The absolute quantities of mRNA copy numbers for each gene were calculated using the qPCR standard curve application provided by the QuantStudio system (Thermo Fisher Scientific).

### 2.5 | *Saccharomyces cerevisiae* transcriptome data acquisition and analysis

A total of 32 *S. cerevisiae* S288C strain transcriptome datasets were retrieved from the NCBI (ncbi.nlm.nih.gov/sra; Accession: PRJNA395936, Gurvich et al., 2017), using the SRA toolkit (function: fastq-dump). The list of transcriptome data downloaded is available in the supplementary information (Supp. Table 2). The downstream analysis of this transcriptome dataset followed the same pipeline as the *D. magna* analysis. The reference genome, gff format file and the GO information terms of *S. cerevisiae* were retrieved from the Saccharomyces Genome Database (yeastgenome.org; Cherry et al., 2012).

### 2.6 | Data visualization

Figures were plotted using the R package ggplot2 (Wickham, 2011) in the R platform. R script for the plots is provided in Zenodo Repository (https://doi.org/10.5281/zenodo.5547206).

## 3 | RESULTS

### 3.1 | Somatic growth rate of *Daphnia magna*

Both temperature and food concentration treatments had significant effects on the somatic growth rate of *D. magna* (Temperature: *F*_2,36_ = 88.71, *P* < 0.01**; Food concentration: *F*_2,36_ = 154.41, *P* < 0.01**; Interaction: *F*_4,36_ = 4.29, *P* < 0.01**; Fig. 1).

**FIGURE 1.**
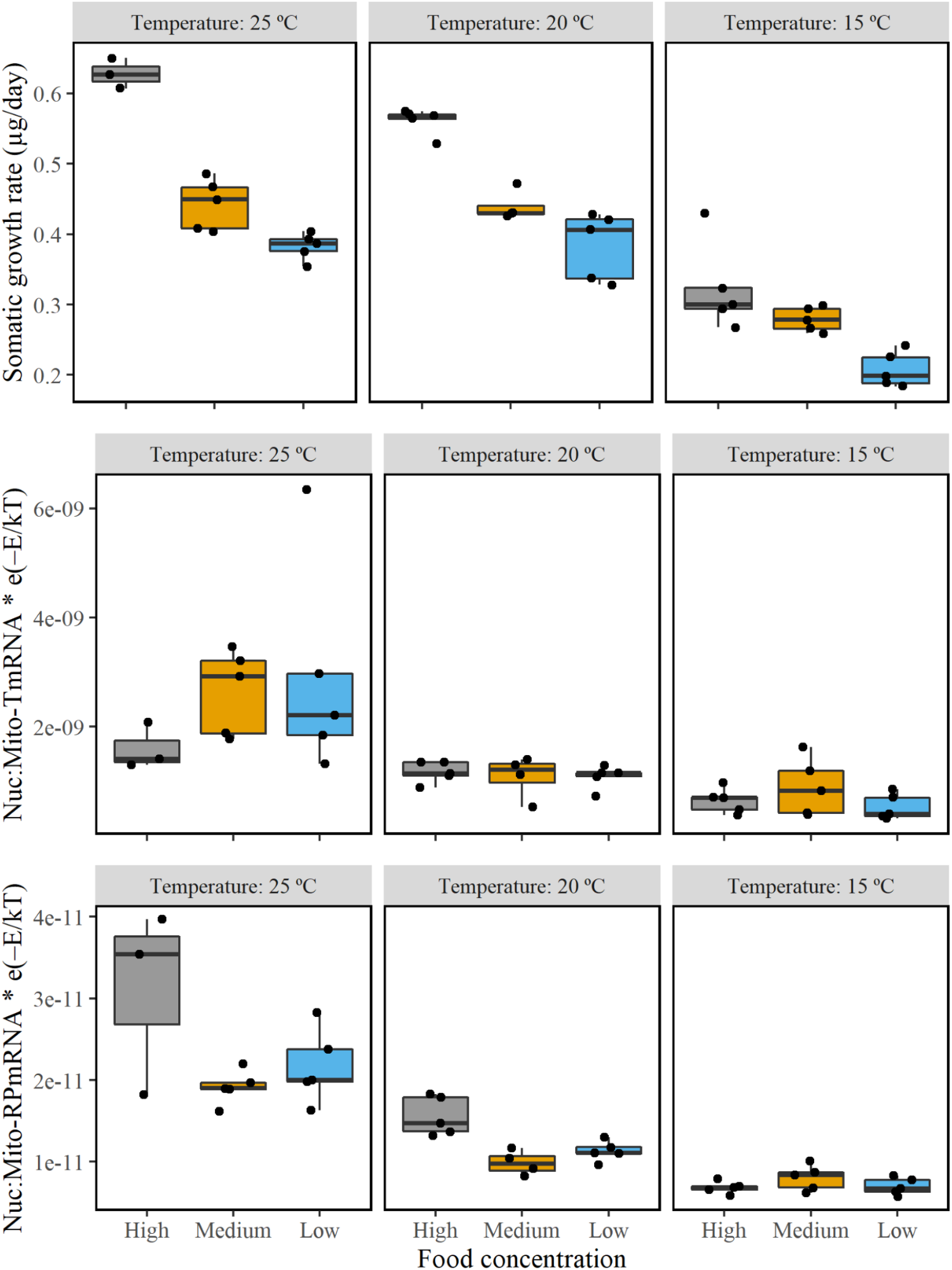
Boxplots on the top show the somatic growth rate (μg/day) of *Daphnia magna* under different temperature and food concentration treatments. The middle boxplot shows the total nuclear and mitochondrial mRNA ratios under different treatments. The bottom boxplot shows the nuclear and mitochondrial ribosomal protein mRNA ratios under different treatments. Both ratios were temperature corrected by the exponential (e) of negative activation energy (–*E*) divided by Boltzmann’s constant (*k*) and temperature in kelvins (*T*).

### 3.2 | *Daphnia magna* transcriptome data output

A total of 45 transcriptome libraries were constructed. Each library yielded about 1,000,000 single-end mRNA reads. One of the libraries had small reads output, therefore removed from the further analysis (a sample treated with high temperature and low food). On average, 87% of mRNA reads were assigned to protein-coding gene regions. The mapping percentages for the libraries were 54%–93% and 2%–16% for the nuclear and the mitochondrial genomes, respectively (Supp. Table 3).

### 3.3 | Relationship between the growth rate and the *Daphnia* transcriptome estimated ratios

#### 3.3.1 | Total nuclear and mitochondrial mRNA ratio

Using Pearson’s correlation test, we found that there is a significant linear correlation between the somatic growth rate of the samples and temperature-corrected Nuc:Mito-TmRNA (*r41* = 0.42, *P* < 0.01*; Fig. 2). Temperature had a significant effect on the total nuclear and mitochondrial mRNA ratio (Nuc:Mito-TmRNA) after temperature correction using the van’t Hoff–Arrhenius relation (*F2*_,34_ = 3.25, *P* < 0.01**), but food concentration and the interaction effect between these two treatments do not show any significant effect on the mRNA ratio (Food concentration: *F*_2,34_ = 2.29, *P* = 0.12; Interaction effect: *F4*_,34_ = 0.77, *P* = 0.55; Fig. 1).

**FIGURE 2.**
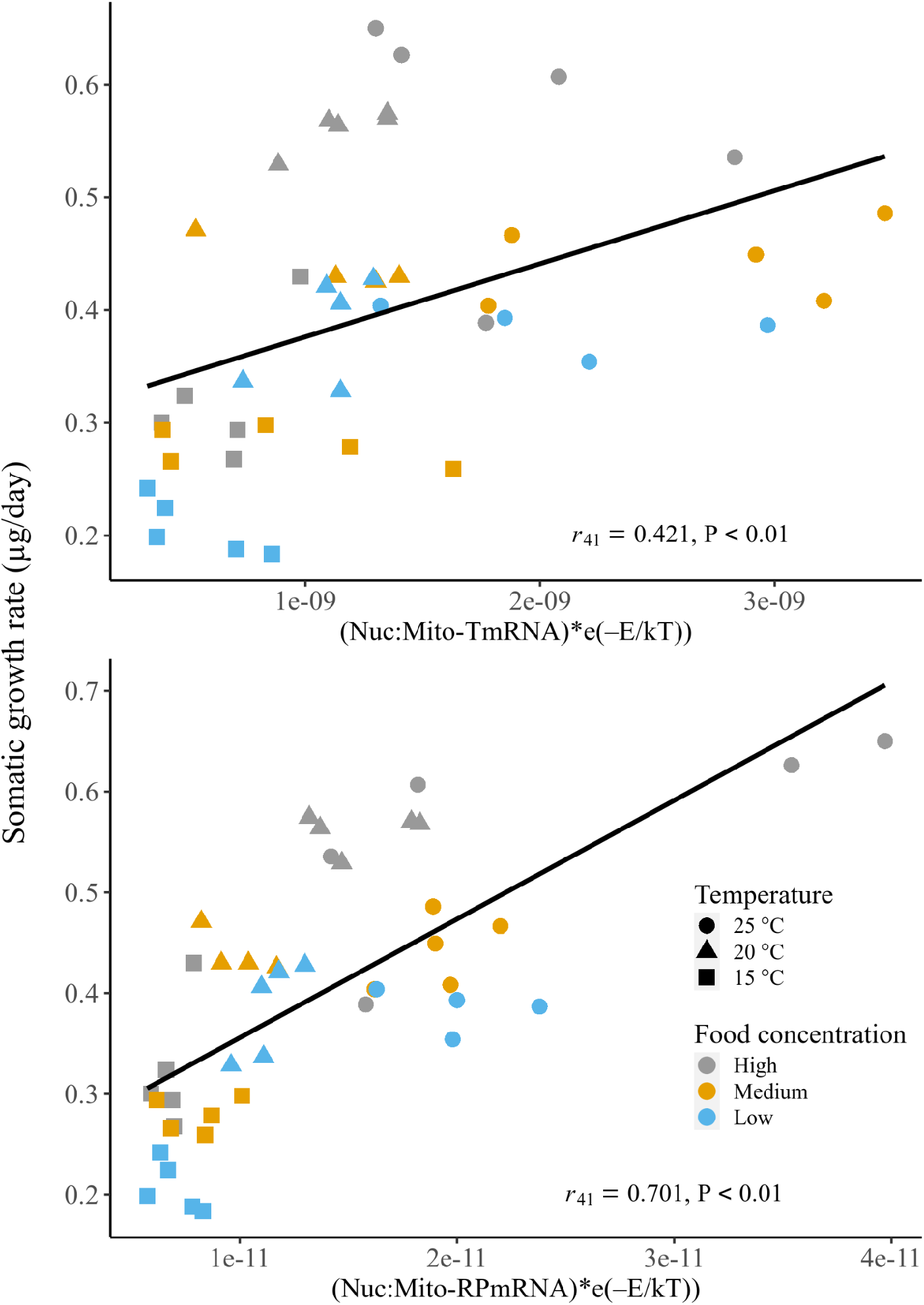
Scatterplots show the total nuclear and mitochondrial mRNA ratio (top) and the nuclear and mitochondrial ribosomal protein mRNA ratio (bottom) against somatic growth rate (μg/day).

#### 3.3.2 | Nuclear and mitochondrial ribosomal protein mRNA ratio

There is a significant linear correlation between the temperature-corrected Nuc:Mito-RPmRNA and somatic growth rate (*r41* = 0.70, *P* < 0.01**; Fig. 2). Both temperature and food concentration treatments had significant effects on the mRNA ratio (Nuc:Mito-RPmRNA), (Temperature: *F2,34* = 37.09, *P* < 0.01**; Food concentration: *F*_2,34_ = 6.38, *P* < 0.01**; Interaction: *F*_4,34_ = 7.00, *P* < 0.01**; Fig. 1). The sample’s biomass had a significant effect on the temperature-corrected Nuc:Mito-RPmRNA (*r*_41_ = 0.52, *P* < 0.01**), and this significant effect can also be observed with the 20 °C treatment (*r*_12_ = 0.78, *P* < 0.01**) and high-food-concentration treatment (*r*_13_ = 0.69, *P* < 0.01**; Fig. 3).

**FIGURE 3.**
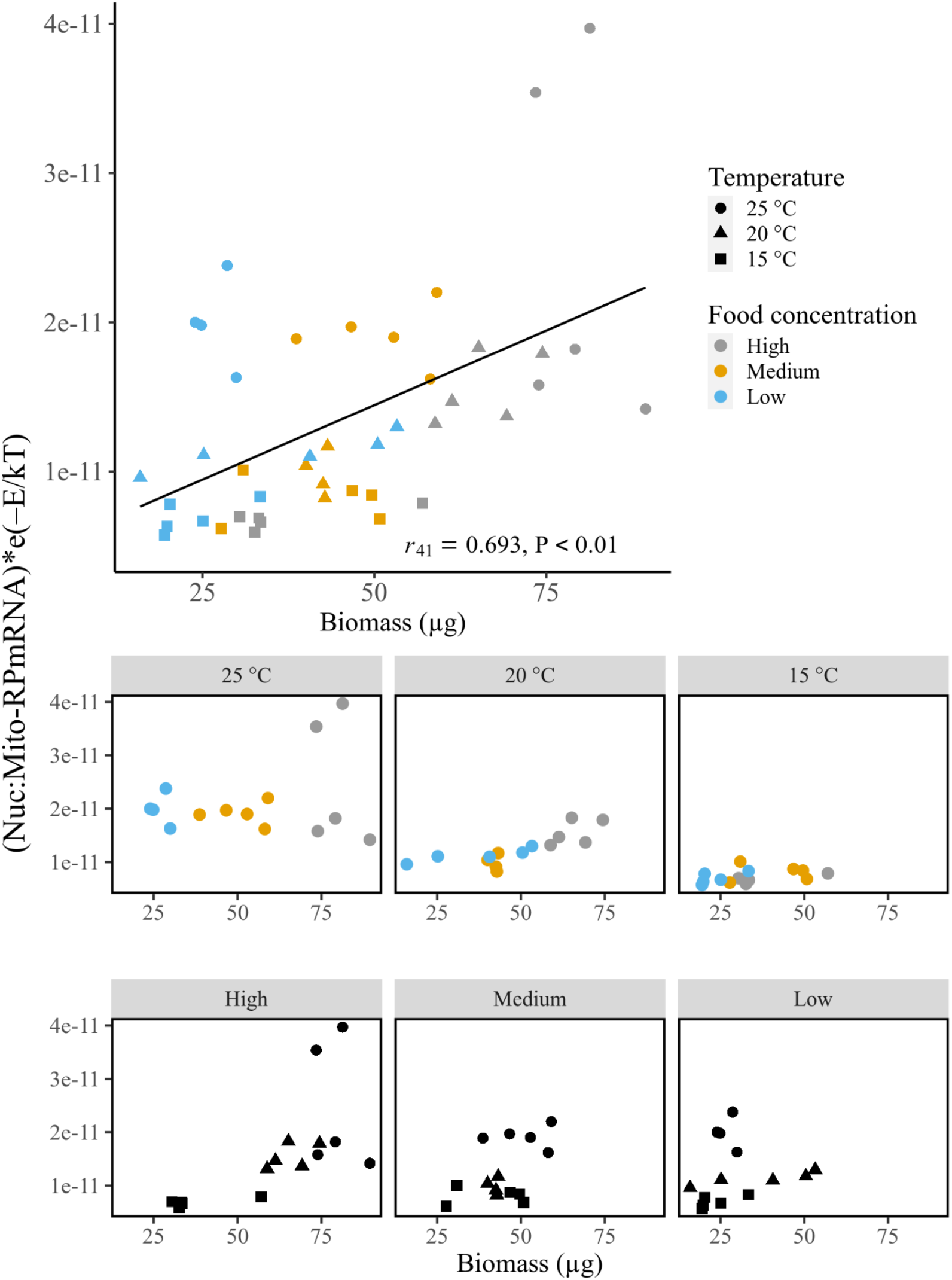
Top scatterplot shows the relationship between the nuclear and mitochondrial ribosomal protein mRNA ratio and biomass. The middle and bottom scatterplots show the relationship between the ratio and temperature and food concentration treatments. High-food-concentration treated samples had a significant correlation.

#### 3.3.3 | Gene ontology terms and the total mitochondrial mRNA ratio

A total of 721 GO terms were used for the ratio estimations for the *D. magna* RNA-Seq dataset. However, the correlation coefficient between the ratios and somatic growth rate were relatively low (<0.49) (Supp. Table 4).

#### 3.3.4 | Nuclear and mitochondrial specific gene mRNA ratio

The mRNA read abundances of a total of 213,240 gene pairs from 26,655 nuclear and eight mitochondrial genes were used to estimate the ratio. Those genes were picked because at least one sequence was mapped for those genes’ reference. Out of all the nuclear and mitochondrial gene pair combinations, the ratios of 31,171 gene pair combinations had a significant correlation with the growth rate. Figure 4 shows the correlation coefficients observed between the nuclear specific and each mitochondrial gene. This result demonstrates that gene pairs with the Cytochrome c oxidase subunit I (COX1) have a better correlation with growth rate than other mitochondrial genes. Therefore, the rest of the calculation will be performed using the mitochondrial COX1 as the denominator.

**FIGURE 4.**
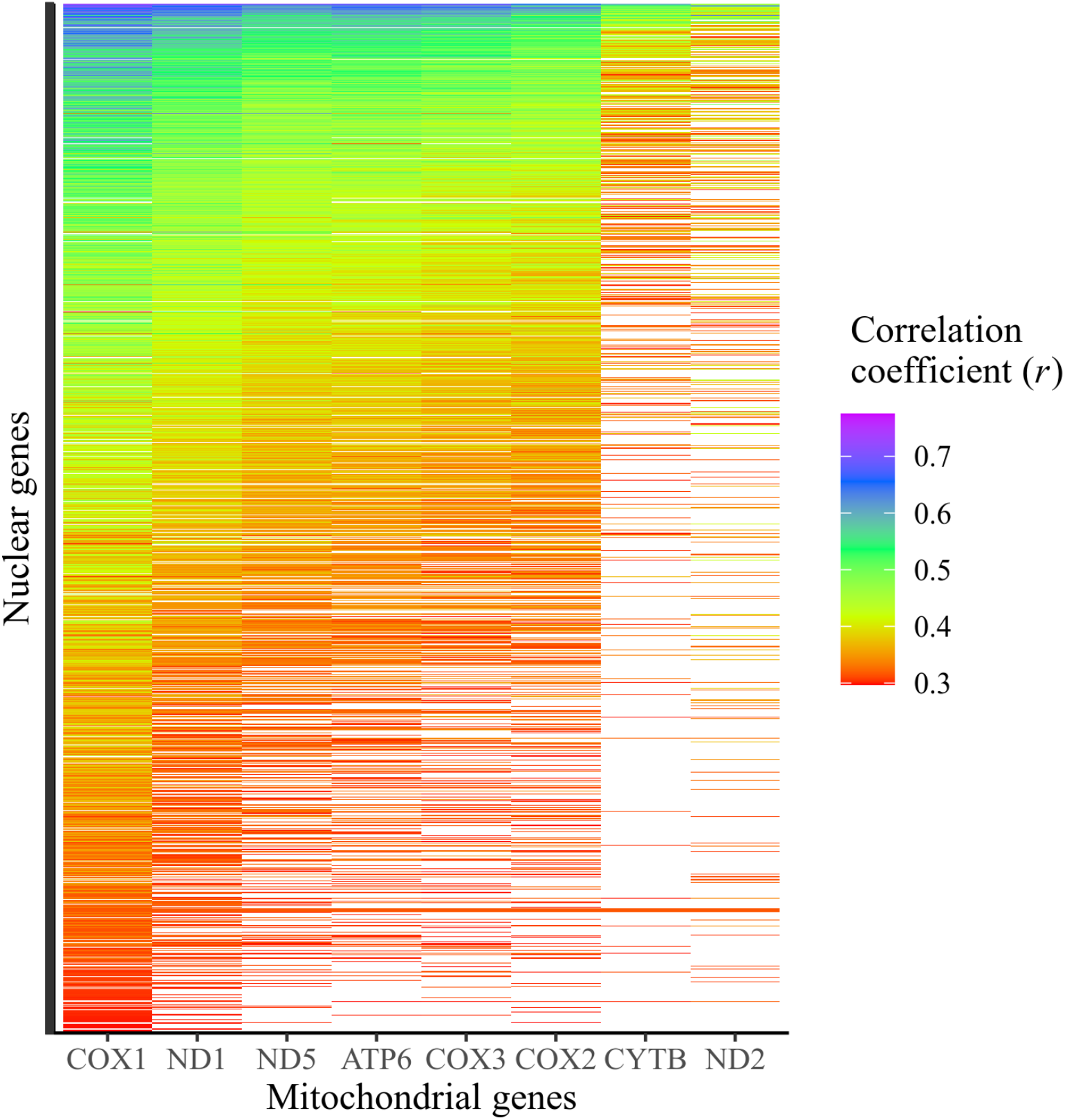
The heatmap shows the correlation coefficient between each nuclear and mitochondrial gene pair combination. Each point of the vertical axis represents a single nuclear gene. Purple colour indicates the gene pair has a high correlation, whereas white colour signifies no significant correlation.

Using the mitochondrial COX1 gene read abundance as the denominator, we extracted the 10 nuclear genes with high correlation coefficient for further examination (Table 1, Fig. 5).

**TABLE 1.**
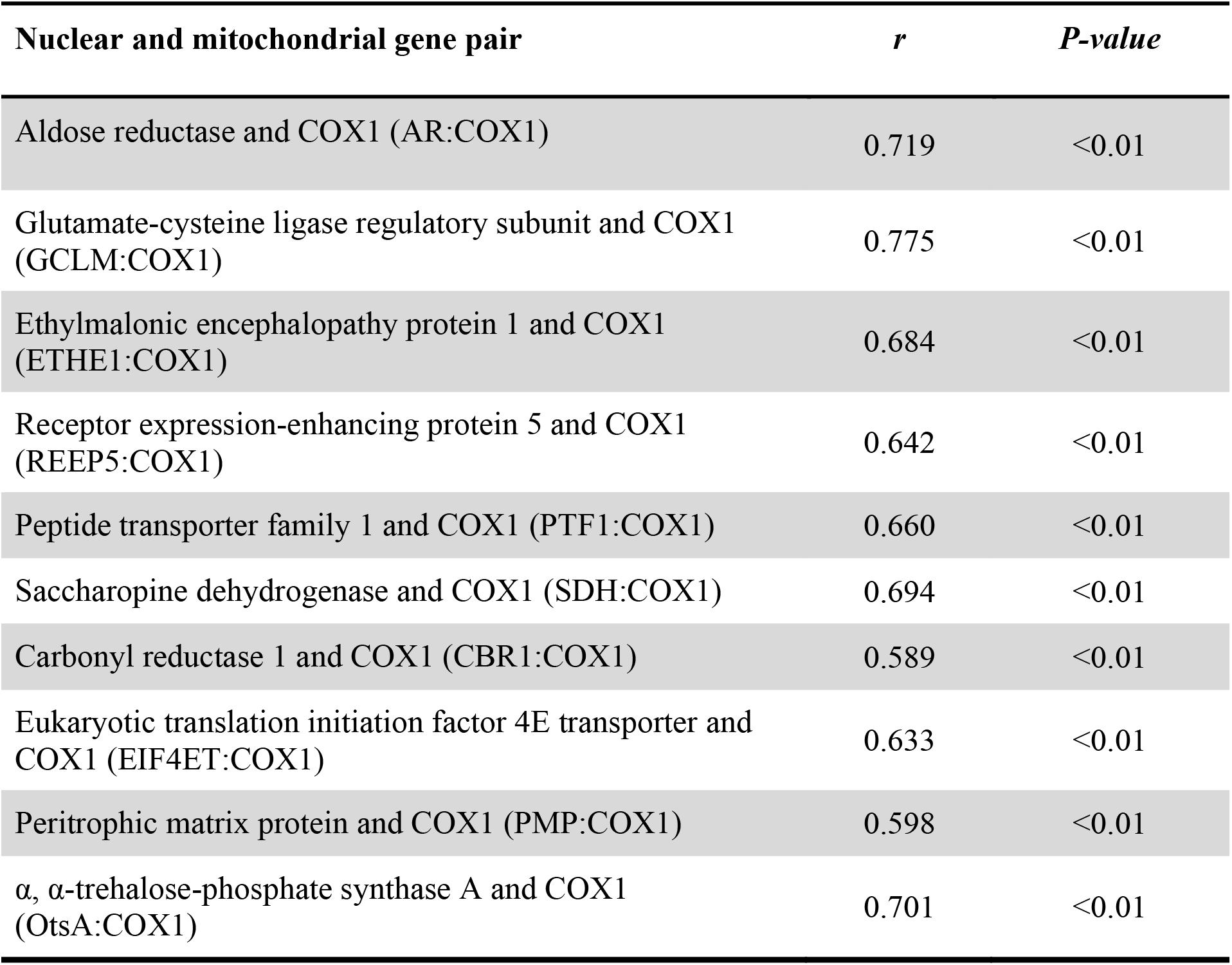
The 10 nuclear and mitochondrial gene pairs that show significant correlation with somatic growth rate in the *D. magna* transcriptome dataset

**FIGURE 5.**
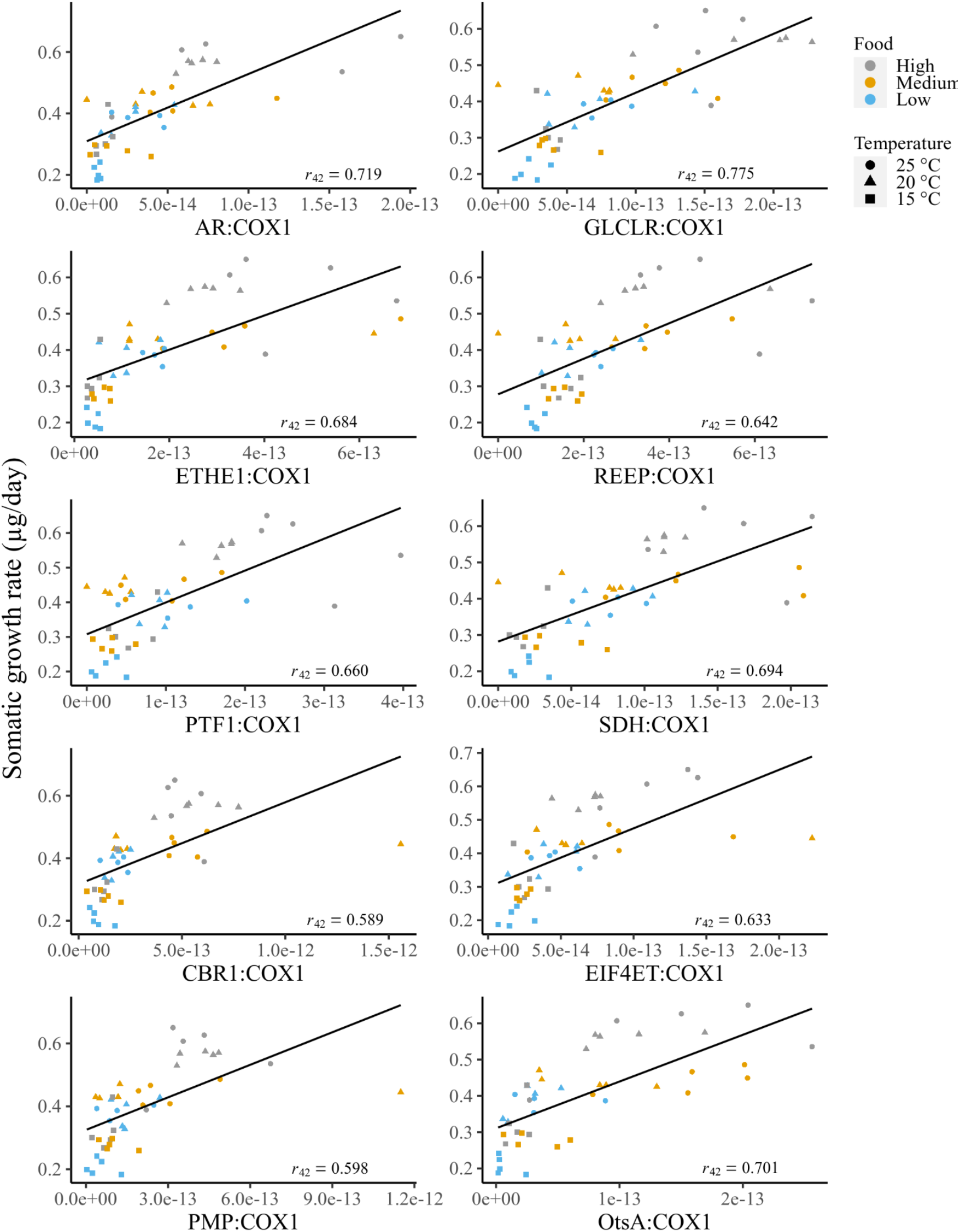
Scatterplots of 10 gene pairs with nuclear and mitochondrial specific gene mRNA ratio that showed a significant correlation with somatic growth rate, using COX1 mRNA abundance as the denominator.

### 3.4 | Validation of nuclear and mitochondrial specific gene mRNA ratios using real-time qPCR

The specific gene mRNA ratios selected from the transcriptome data were validated using absolute real-time qPCR. We designed multiple primers for the 10 nuclear genes. However, only six genes: aldose reductase (AR), glutamate-cysteine ligase regulatory subunit (GCLM), ethylmalonic encephalopathy protein 1 (ETHE1), receptor expression-enhancing protein 5 (REEP5), peptide transporter family 1 (PTF1), saccharopine dehydrogenase (SDH), carbonyl reductase (CBR1), eukaryotic translation initial factor 4E transporter (EIF4ET), peritrophic matrix protein (PMP) and α, α-trehalose-phosphate synthase A (OtsA), were successfully amplified due to the strict requirements for the real-time qPCR experiment (Taylor et al., 2019). The average cDNA copy numbers for each gene that was detected and quantified using the realtime qPCR are 1,428,089, 17, 2, 6, 79, 13 and 6,494 copies for COX1, REEP5, PTF1, CBR1, OtsA, AR and ETHE1, respectively. The ratios with OtsA, AR and ETHE1 showed significant correlation with somatic growth rate (OtsA: *n* = 27, *r* = 0.854, *P* < 0.01**; ETHE1: *n* = 19, *r* = 0.761, *P* < 0.01**; AR: *n* = 27, *r* = 0.702, *P* < 0.01**; Fig. 6).

**FIGURE 6.**
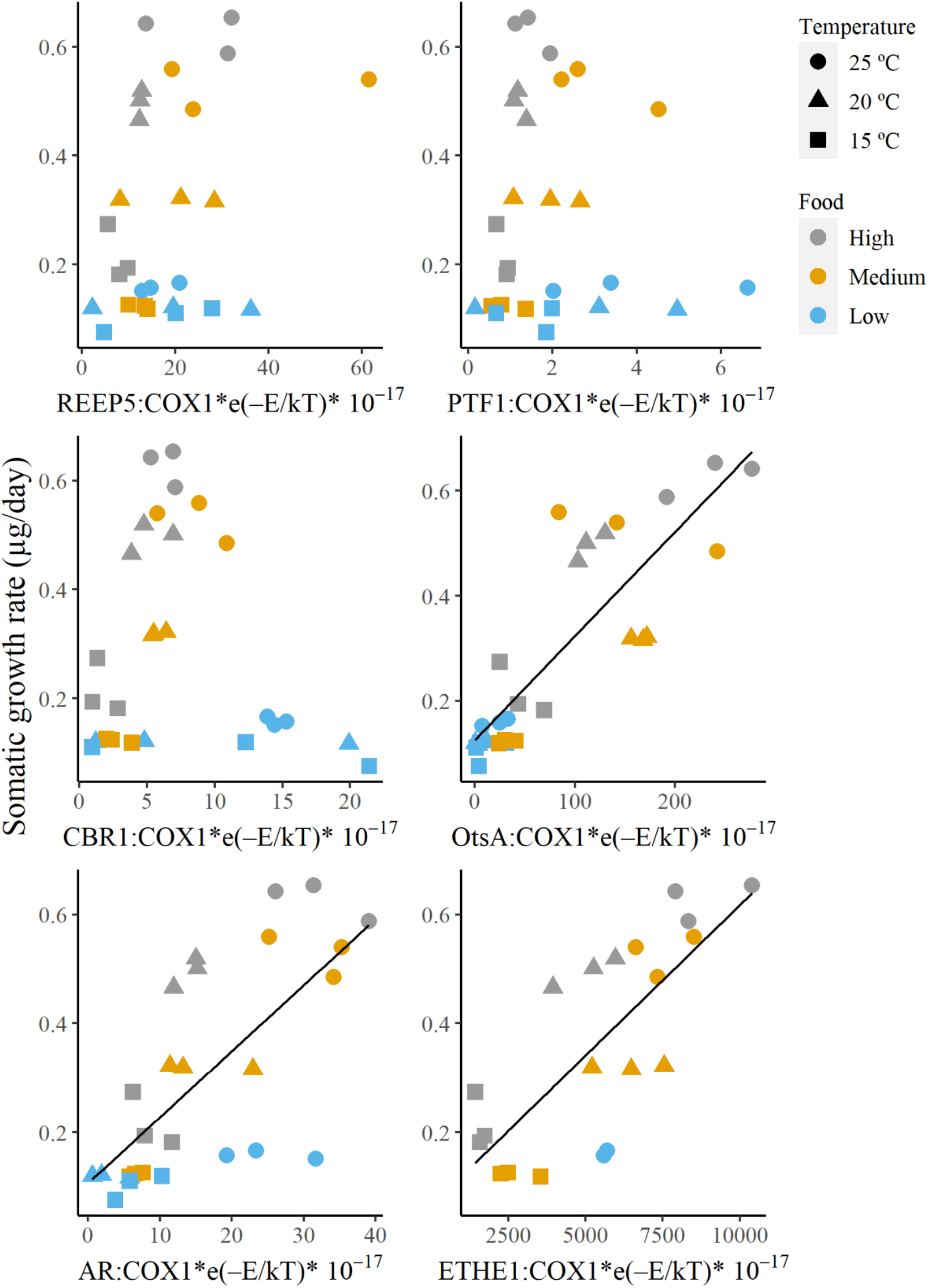
Scatterplots of nuclear and mitochondrial specific gene mRNA ratio against somatic growth rate (top left: REEP5, top right: PTF1, middle left: CBR1, middle right: OtsA, bottom left: AR and bottom right: ETHE1). The ratios were estimated using real-time qPCR.

### 3.5 | Messenger RNA ratios using the *Saccharomyces cerevisiae* transcriptome dataset

Pearson’s correlation test showed that both Nuc:Mito-TmRNA and Nuc:Mito-RPmRNA ratios correlated significantly with growth rate for *S. cerevisiae* (Nuc:Mito-TmRNA: *r*_26_ = 0.785, *P* < 0.01**; Nuc:Mito-RPmRNA: *r*_26_ = 0.823, *P* < 0.01**; Fig. 7). The analysis of variance test showed that phosphorus levels have a significant effect on both Nuc:Mito-TmRNA (*F*_3,24_ = 5.83, *P* < 0.01**) and Nuc:Mito-RPmRNA (*F*_3,24_ = 4.42, *P* = 0.01*). The Tukey ad hoc test showed that the mRNA ratios from the no-phosphorus treatment group have significant differences from other groups. However, the correlation is only significant in the low-phosphorus-level groups, namely, *S. cerevisiae* that are treated with 0 mM and 0.06 mM phosphorus (0 mM: *r*_5_ = 0.947, *P* < 0.01**; 0.06 mM: *r*_5_ = 0.846, *P* = 0.02*) for Nuc:Mito-TmRNA. For Nuc:Mito-RPmRNA, only the 0.06 mM phosphorus-level treatment group had a significant correlation between the mRNA ratio and growth rate (*r*_5_ = 0.916, *P* < 0.01**).

**FIGURE 7.**
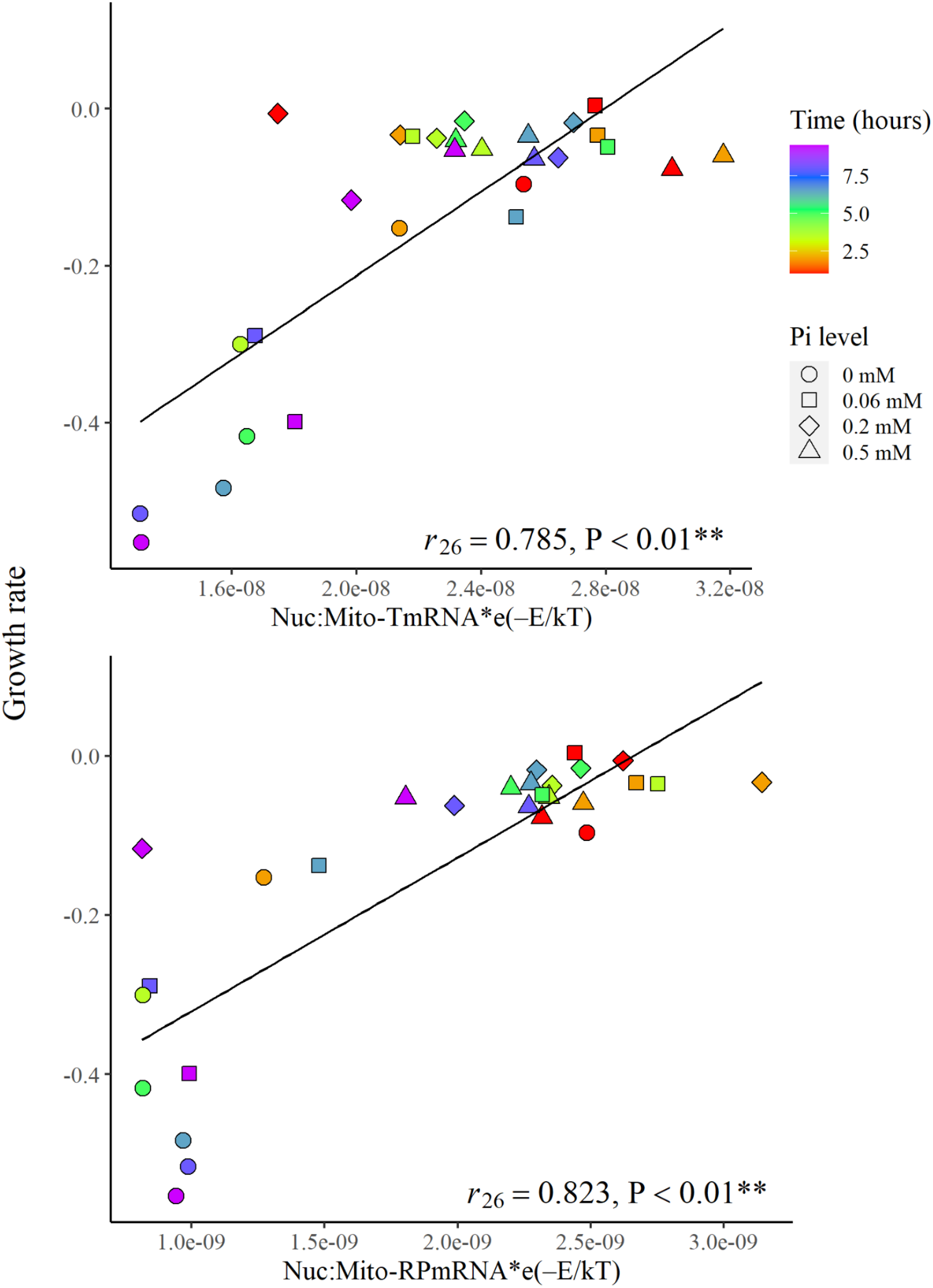
The scatterplots show the relationship between the ratios and the growth rate of *S. cerevisiae*. The top and bottom scatterplots show the relationships of the total and ribosomal protein mRNA ratios, respectively.

A large number of GO term ratios (each GO term read number and the total mitochondrial mRNA) had significant correlations with the growth rate. The GO term with the highest correlation coefficient is the cytoplasmic translation (GO: 0002181). Other GO terms, including organelle assembly (GO: 0070925), ribosomal small subunit biogenesis (GO: 0042274), ribosome assembly (GO: 0042255) and ribosomal large subunit biogenesis (GO: 0042273), also correlated with growth rate. Table 2 lists the top 10 GO terms that correlate with the samples’ growth rate.

**TABLE 2.** The temperature-corrected GO mRNA ratio of the 10 listed GO terms that showed the highest correlation coefficient (*r*) with the growth rate of the *S. cerevisiae* sample

| GO term | $r$ | $P$-value |
| --- | --- | --- |
| Cytoplasmic translation (GO: 0002181) | 0.887 | <0.01 |
| Nucleobase-containing compound transport (GO: 0015931) | 0.884 | <0.01 |
| Organelle assembly (GO: 0070925) | 0.880 | <0.01 |
| Ribosomal small subunit biogenesis (GO:0042274) | 0.874 | <0.01 |
| RNA splicing (GO: 0008380) | 0.871 | <0.01 |
| Ribosome assembly (GO: 0042255) | 0.867 | <0.01 |
| rRNA processing (GO: 0006364) | 0.863 | <0.01 |
| Translational elongation (GO: 0006414) | 0.860 | <0.01 |
| Ribosomal large subunit biogenesis (GO: 0042273) | 0.850 | <0.01 |
| Ribosomal subunit export from nucleus (GO: 0000054) | 0.835 | <0.01 |

## 4 | DISCUSSION

According to endosymbiotic theory, the origin of mitochondria was a free-roaming nucleus-bearing cell that was engulfed by another cell and gradually evolved into the current state of mitochondria in eukaryotic cells (Margulis & Bermudes, 1985). During the evolutionary process, fragments of mitochondrial genomes migrated into the cell nucleus and integrated into the nuclear genome (Lopez et al., 1994). The migration resulted in the compacts of mitochondrial genome size, and the remaining genes are mainly involved in energy production for cellular processes (Adams & Palmer, 2003). Thus, the rest of the biological processes such as cell growth and division rely on the genes expressed in the nuclear genome. All of the mRNA-based growth indices proposed in this study use these function differences as the ratios.

### 4.1 | Nuc:Mito-TmRNA and Nuc:Mito-RPmRNA as the growth rate indices

Both Nuc:Mito-TmRNA and Nuc:Mito-RPmRNA showed significant correlations with the growth rate of *D. magna* and *S. cerevisiae*. We also found that temperature and food concentration treatments significantly affect Nuc:Mito-RPmRNA ratios, whereas the Nuc:Mito-TmRNA ratio is only significantly affected by temperature in *D. magna*. This might be one of the reasons for the weaker correlation of Nuc:Mito-TmRNA with somatic growth rate compared with Nuc:Mito-RPmRNA. The rationales of those two indices are very similar: separating cell growth and division (measured by nuclear-encoded gene expression) and energy production (measured by the mitochondrial gene expression) components from the total mRNA abundance and using them as the growth indices. The difference between the two ratios is that the Nuc:Mito-TmRNA includes every protein-coding gene of nuclear and mitochondrial genomes, whereas the Nuc:Mito-RPmRNA only considers the projected abundance of ribosome molecules. The synthesis of ribosomes requires a large amount of environmentally limited nutrients such as nitrogen and phosphorus (Warner, 1999). Hence, the change of Nuc:Mito-RPmRNA is expected to be slow. In contrast, less energy and nutrients are required for the dynamics of the Nuc:Mito-TmRNA. Because of these differences, we expect that Nuc:Mito-TmRNA has higher sensitivity than Nuc:Mito-RPmRNA. Despite the large taxonomic distance, Nuc:Mito-TmRNA and Nuc:Mito-RPmRNA of both *D. magna* and *S. cerevisiae* showed significant correlations against growth rate. This observation indicates that the two mRNA ratio indices have the potential to be universal growth rate indices because all of the Eukaryota [except several species of parasitic amoebozoa that consist of mitosome instead of mitochondria (Tovar et al., 1999)] rely on the mitochondria-related genes to synthesize proteins for energy generation, whereas the nuclear genes produce the rest of the proteins that contribute to cell growth and divisions.

### 4.3 | GO terms and total mitochondrial mRNA ratio

There are many GO terms in the biological process domain that are directly or indirectly involved in growth. For example, the genes that are annotated to cytoplasmic translation (GO: 0002181) are responsible for the protein translation process in the cytoplasm. In the *S. cerevisiae* dataset, the mRNA ratio using this GO term showed the strongest correlation with growth rate. The uniqueness of the GO terms and mitochondrial mRNA ratio is that they can minimize the gene expression noise caused by other genes that do not contribute to the organism’s growth, such as secondary metabolites. However, we did not observe similar patterns in the *D. magna* transcriptome dataset. The results of our *D. magna* dataset showed that only 11.4% (3,052 out of 26,655 genes) of the protein-coding genes were annotated to the GO terms, whereas the estimate for *S. cerevisiae* was 89.2% (5,882 out of 6,572 genes). This result indicates that the majority of the genes of *D. magna* (88.6%) do not have a known function or require thorough functional annotation to assign proper GO terms. This suggests that a thorough transcriptome functional annotation is required for this mRNA ratio index, which indicates that the GO terms and total mitochondrial mRNA ratio might be less effective for species that lack information on the annotated genes’ function.

### 4.4 | Specific nuclear gene and COX1 mRNA ratio

The correlation coefficients estimated from the mitochondrial COX1 gene were the best among the mitochondrial genes (Fig. 4). This result demonstrated that the mitochondrial COX1 was the best gene to be used as the denominator.

First, we estimated candidate nuclear genes from the transcriptome dataset. Next, we validated the combination with real-time qPCR. Our results demonstrated that three pairs of nuclear and mitochondrial gene mRNA ratios, namely, OtsA:COX1-mRNA, AR:COX1-mRNA and ETHE1:COX1-mRNA, are good candidates for estimating growth rate due to consistent results in both transcriptome and real-time qPCR. In the present study, we focused only on the three genes. However, it is expected that more genes can be found with more extensive datasets.

### 4.5 | Advantages and disadvantages of mRNA ratios as the growth rate indices

This is the first study that proposed growth rate indices involving protein-coding gene mRNA read abundance from transcriptome high-throughput sequencing data. The four types of proposed nuclear and mitochondrial mRNA ratios each come with their advantages and disadvantages (Table 3). First, Nuc:Mito-TmRNA can be calculated by aligning transcriptome mRNA reads to a mitochondrial reference, whereas the rest of the reads will be the nuclear mRNA reads. Hence, this method can be utilized by non-model species due to the availability of mitochondrial reference databases (Leray et al., 2018; Sato et al., 2018). However, this approach includes all protein-coding genes, some of which might not contribute to growth. Besides, this approach requires high-throughput transcriptome sequencing technology.

**TABLE 3.** List of advantages and disadvantages of each proposed mRNA ratio index

| Advantages | Disadvantages |
| --- | --- |
| **Total nuclear and mitochondrial mRNA ratio (Nuc:Mito-TmRNA)** |  |
| • Requires only the mitochondrial genome databases to perform the calculation.<br>• Applicable across different taxonomic groups. | • Lacks specificity—all protein-coding genes are included in the calculation.<br>• Requires high-throughput sequencing technology. |
| **Nuclear and mitochondrial ribosomal protein mRNA ratio (Nuc:Mito-RPmRNA)** |  |
| • Applicable across different taxonomic groups. | • Requires both cytosolic and mitochondrial ribosomal protein sequence databases.<br>• Requires high-throughput sequencing technology. |
| **GO term and total mitochondrial mRNA ratio** |  |
| • Specific on the function of the genes included—potentially less noise. | • Requires GO annotation.<br>• Requires high-throughput sequencing technology. |
| **Nuclear and mitochondrial specific gene mRNA ratio** |  |
| • Can be conducted through the qPCR platform. | • Homolog could not be found in some species for certain genes. |

For the nuclear and mitochondrial ribosomal protein mRNA ratio (Nuc:Mito-RPmRNA), the main advantage is the high potential of this mRNA ratio in predicting the somatic growth rate due to the high correlation coefficient between these two variables. A reference database of ribosomal protein is also available (Nakao et al., 2004), but the species coverage is not as widely available as the mitochondrial reference genome database. Similar to Nuc:Mito-TmRNA, the Nuc:Mit-RPmRNA approach also requires a high-throughput sequencing platform.

Compared with Nuc:Mito-TmRNA and Nuc:Mito-RPmRNA, the GO terms and total mitochondrial mRNA ratio provide higher specificity, which minimizes potential noise generated by protein-coding genes, which is not related to growth. However, the method requires a well-established GO terms reference. Currently, such information is mainly available only for the major model organisms. A high-throughput sequencing platform is also needed for this approach.

Lastly, the nuclear and mitochondrial specific gene mRNA ratio has a unique advantage because this approach can be performed using the real-time qPCR platform. The method also has very high sensitivity. However, not all species have the homolog genes of the nuclear gene proposed in this study, such as the homolog of ETHE1, which is yet to be discovered in *S. cerevisiae*.

Cautions are required when the ratios are applied for non-metazoan eukaryote species. Enrichment of mRNA using polyadenylation (Oligo dT purification) is the standard strategy in transcriptome library preparations. However, the polyadenylation of non-metazoan eukaryote mitochondrial mRNA is known to promote degradation (Chang & Tong, 2012). Therefore, it is likely that the data generated from the Oligo dT-enriched libraries will not provide good mitochondrial mRNA abundance estimates. Instead, other mRNA enrichment, such as ribosomal RNA depletion, is recommended.

### 4.6 | Prospect of the proposed mRNA indices with metatranscriptomics

Metatranscriptomics analysis has gained popularity in the past few years. Aside from the microbial community, eukaryotes such as zooplanktons have also pursued the same approach recently (Damon et al., 2012; Lenz et al., 2021; Lopez et al., in press; Machida et al., 2021). All of the ratios proposed in this study can be estimated from metatranscriptome datasets. Therefore, we can estimate diversity indices, species composition and growth indices from a single metatranscriptome sequence dataset. Reference sequences of some of the genes proposed in this study are less available. Therefore, deposition of those functional gene sequences will be required before fully utilizing some of the methods.

## ACKNOWLEDGEMENTS

The first author thanks his thesis committee members for providing helpful advice in preparing the manuscript. We also thank the NGS High Throughput Genomics Core at the Biodiversity Research Center. W.L.K. was supported by the Taiwan International Graduate Program scholarship for his PhD degree. This project was supported by Academia Sinica, Taiwan (R.J.M.), the Ministry of Science and Technology, Taiwan (108-2611-M-001, 109-2611-M-001, 110-2611-M-001; R.J.M.) and the Scientific Committee on Oceanic Research working group 157 (R.J.M.). The funding agencies played no part in the study design, data collection, analysis, decision to publish or manuscript preparation.

## AUTHOR CONTRIBUTIONS

The experiment was conceived by R.J.M. and W.L.K.; Animal culturing and molecular experiments were performed by W.L.K.; Statistical analyses were performed by W.L.K. with guidance from R.J.M.; W.L.K. wrote most of the manuscript with help from R.J.M. R.J.M. edited and approved the final version of the manuscript.

## DATA AVAILABILITY

The RNA-Seq data of *D. magna* was deposited in the Sequence Read Archive section of the National Center for Biological Information (accession number: PRJNA762666). The R script for mRNA ratios calculation (https://doi.org/10.5281/zenodo.5506660) and plots (https://doi.org/10.5281/zenodo.5547206) was uploaded to the Zenodo Repository.

