## Supplemental materials for "Development of transcriptomics-based eukaryotes growth rate indices"

**Supp. Table 1** List of primers used in real-time qPCR. The 'Dmag_pre-' prefix indicates the precursor primers that are used to sequence longer products for standard curve preparation for absolute quantification.

| Primer | Size (bp) | Sequence (5′ to 3′) | GC (%) | $T_m$ (°C) |
|---|---|---|---|---|
| Dmag_preCOX1_1377F | 18 | GGG TTT GAT CAG GTA TGG | 50 | 48.0 |
| Dmag_preCOX1_2502R | 18 | GGA TAC CAA TGA GCA ACC | 50 | 48.0 |
| DmagCOX1_159F | 20 | ACC TCC TGC ACT AAC TCT TT | 45 | 49.7 |
| DmagCOX1_259R | 20 | CAT GAG CAA TTC CAG CAG AT | 45 | 49.7 |
| Dmag_preREEP5_65F | 20 | GTT GCT CGC TTC ACT GTT GG | 55 | 53.8 |
| Dmag_preREEP5_948R | 20 | GCC AAC CAG TAC AAG GGG AA | 55 | 53.8 |
| DmagREEP5_18F | 20 | CAC GTC TCT ACC AAC GGC TC | 60 | 55.9 |
| DmagREEP5_195R | 19 | GAA GAA ACG GTT GGG CAG C | 58 | 53.2 |
| Dmag_prePTF1_251F | 20 | TCG GAG GAG GAA GAA GAC GA | 55 | 53.8 |
| Dmag_prePTF1_1099R | 20 | GCA ACG ACA CCG GAA GTA GA | 55 | 53.8 |
| DmagPTF1_299F | 20 | CAG CTA GTC CTA CCT GAG AA | 50 | 51.8 |
| DmagPTF1_400R | 20 | ACA TCC TCT GGT TAC TGC CT | 50 | 51.8 |
| Dmag_preCBR1_1379F | 20 | GCC GCT GGT AGC CTA ATC AT | 55 | 53.8 |
| Dmag_preCBR1_2306R | 20 | CTC CAC CCC AAC ATT CAC CA | 55 | 53.8 |
| DmagCBR1_154F | 20 | TGG TGT TTC GCA GAG AAC CC | 55 | 53.8 |
| DmagCBR1_242R | 20 | TCA CAA CAG ACG GAC AGG GA | 55 | 53.8 |
| Dmag_preOtsA_587F | 20 | TCC AAT CCA CTC CGA AAG CC | 55 | 53.8 |
| Dmag_preOtsA_1896R | 20 | GGG TGT CGA TCG TTT GGA CT | 55 | 53.8 |

| | | | | |
|---|---|---|---|---|
| DmagOtsA_351F | 20 | GGC AAC CAT TGA CTT GAC GC | 55 | 53.8 |
| DmagOtsA_450R | 20 | CGA AAC TAA ACG CGT CCT GG | 55 | 53.8 |
| Dmag_preAR_1432F | 20 | GAC ATT GTG GTG ACC GCC TA | 55 | 53.8 |
| Dmag_preAR_2510R | 20 | CAG TCG CTT CCT GCT GTA CA | 55 | 53.8 |
| DmagAR_387F | 20 | GGT GGT GGT GGT AAA CAT CC | 55 | 53.8 |
| DmagAR_486R | 20 | GCT ACA ACG AAA CCC GAC TC | 55 | 53.8 |
| Dmag_preETHE1_364F | 18 | CCT ACG TTT GCC ATG AAC | 50 | 48.0 |
| Dmag_preETHE1_500R | 18 | TGG CAG GGA GAA GAT ATG | 50 | 48.0 |
| DmagETHE1_148F | 18 | TGT GGT CGC ACC GAT TTT | 50 | 48.0 |
| DmagETHE1_293R | 19 | TTC TCC TCC CAT ACA GTC G | 53 | 51.1 |

*GC(%) - guanine-cytosine content
Tm(°C) - Melting temperature

**Supp. Table 2** List of transcriptome datasets with their accession numbers for each experiment run downloaded from SRA (Accession: PRJNA395936).

| Accession: experiment runs | Phosphorus level (mM) | Time (hours) |
|---|---|---|
| SRR5872941 | 0.20 | 9.5 |
| SRR5872942 | 0.20 | 8.0 |
| SRR5872943 | 0.20 | 6.5 |
| SRR5872944 | 0.20 | 5.0 |
| SRR5872945 | 0.20 | 3.5 |
| SRR5872946 | 0.20 | 2.0 |
| SRR5873005 | 0.06 | 24.7 |
| SRR5873006 | 0.06 | 1.0 |
| SRR5873007 | 0.06 | 3.5 |
| SRR5873008 | 0.06 | 2.0 |
| SRR5873009 | 0.06 | 6.5 |
| SRR5873010 | 0.06 | 5.0 |
| SRR5873011 | 0.06 | 9.5 |

| | | |
|---|---|---|
| SRR5873012 | 0.06 | 8.0 |
| SRR5873034 | 0.00 | 5.0 |
| SRR5873035 | 0.00 | 6.5 |
| SRR5873036 | 0.00 | 8.0 |
| SRR5873037 | 0.00 | 9.5 |
| SRR5873038 | 0.00 | 1.0 |
| SRR5873039 | 0.00 | 24.7 |
| SRR5873040 | 0.00 | 2.0 |
| SRR5873041 | 0.00 | 3.5 |
| SRR5873076 | 0.50 | 5.0 |
| SRR5873077 | 0.50 | 6.5 |
| SRR5873078 | 0.50 | 2.0 |
| SRR5873079 | 0.50 | 3.5 |
| SRR5873080 | 0.50 | 1.0 |
| SRR5873081 | 0.50 | 24.67 |
| SRR5873104 | 0.50 | 9.5 |
| SRR5873105 | 0.50 | 8.0 |
| SRR5873119 | 0.20 | 1.0 |

**Supp. Table 3** The mapping results of each RNA-Seq library.

| Sample | Raw read abundance | Mapped mRNA read abundance | Mitochondrial read abundance | Mapping percentage | Mitochondrial mRNA read percentage |
|---|---|---|---|---|---|
| HTHF2 | 969171 | 853639 | 44581 | 88.08% | 5.22% |
| HTHF5 | 721259 | 621704 | 70441 | 86.20% | 11.33% |
| HTHF7 | 1044183 | 925974 | 68264 | 88.68% | 7.37% |
| HTHF8 | 813799 | 694146 | 97473 | 85.30% | 14.04% |
| HTHF10 | 670653 | 570938 | 73730 | 85.13% | 12.91% |
| HTMF2 | 1090580 | 964642 | 75341 | 88.45% | 7.81% |

| | | | | | |
|---|---|---|---|---|---|
| HTMF3 | 985140 | 909308 | 40891 | 92.30% | 4.50% |
| HTMF4 | 1000432 | 920790 | 42946 | 92.04% | 4.66% |
| HTMF8 | 1091692 | 998307 | 56282 | 91.45% | 5.64% |
| HTMF10 | 1022831 | 921496 | 76056 | 90.09% | 8.25% |
| HTLF11 | 900512 | 813005 | 29332 | 90.28% | 3.61% |
| HTLF12 | 1878953 | 1677463 | 107280 | 89.28% | 6.40% |
| HTLF13 | 813711 | 706645 | 78210 | 86.84% | 11.07% |
| HTLF15 | 893250 | 797352 | 54726 | 89.26% | 6.86% |
| MTHF1 | 891381 | 767472 | 54461 | 86.10% | 7.10% |
| MTHF2 | 661593 | 574005 | 72716 | 86.76% | 12.67% |
| MTHF3 | 1000825 | 894630 | 80120 | 89.39% | 8.96% |
| MTHF6 | 946253 | 843516 | 54547 | 89.14% | 6.47% |
| MTHF7 | 1039752 | 900565 | 102494 | 86.61% | 11.38% |
| MTMF1 | 369557 | 198525 | 9610 | 53.72% | 4.84% |
| MTMF3 | 1069284 | 949925 | 78143 | 88.84% | 8.23% |
| MTMF7 | 1370675 | 1230382 | 69997 | 89.76% | 5.69% |
| MTMF8 | 1583977 | 1420494 | 88070 | 89.68% | 6.20% |
| MTMF10 | 1093068 | 918151 | 130287 | 84.00% | 14.19% |
| MTLF36 | 859312 | 732788 | 80650 | 85.28% | 11.01% |
| MTLF37 | 1352868 | 1183414 | 144773 | 87.47% | 12.23% |
| MTLF38 | 1551550 | 1397704 | 115458 | 90.08% | 8.26% |
| MTLF39 | 1072897 | 958619 | 77242 | 89.35% | 8.06% |
| MTLF40 | 1156604 | 1033226 | 76051 | 89.33% | 7.36% |
| LTHF11 | 1211518 | 1071503 | 116919 | 88.44% | 10.91% |
| LTHF13 | 937260 | 814715 | 119196 | 86.93% | 14.63% |
| LTHF14 | 1750669 | 1588096 | 87020 | 90.71% | 5.48% |
| LTHF16 | 1114172 | 998495 | 59322 | 89.62% | 5.94% |
| LTHF17 | 1131142 | 935272 | 140260 | 82.68% | 15.00% |
| LTMF1 | 1016048 | 836370 | 45981 | 82.32% | 5.50% |

| | | | | | |
|---|---|---|---|---|---|
| LTMF2 | 991330 | 858128 | 103124 | 86.56% | 12.02% |
| LTMF3 | 1174518 | 1090338 | 26313 | 92.83% | 2.41% |
| LTMF4 | 1284143 | 1105586 | 156421 | 86.10% | 14.15% |
| LTMF5 | 971392 | 887895 | 35055 | 91.40% | 3.95% |
| LTLF3 | 998673 | 828431 | 133843 | 82.95% | 16.16% |
| LTLF4 | 994554 | 844160 | 108226 | 84.88% | 12.82% |
| LTLF11 | 1172438 | 1002503 | 154165 | 85.51% | 15.38% |
| LTLF13 | 1209253 | 1018971 | 69519 | 84.26% | 6.82% |
| LTLF15 | 1633369 | 1472750 | 93534 | 90.17% | 6.35% |

**Supp. Table 4** The GO terms and total mitochondrial mRNA ratio of the 10 GO terms showing the highest correlation coefficient (*r*) with the somatic growth rate.

| **GO term** | ***r*** | ***P-value*** |
|---|---|---|
| Gama-tubulin complex localization (GO: 0033566) | 0.489 | <0.01 |
| DNA replication, removal of RNA primer (GO: 0043137) | 0.432 | <0.01 |
| Uracil salvage (GO: 0006223) | 0.422 | <0.01 |
| UMP salvage (GO: 0044206) | 0.422 | <0.01 |
| Cell volume homeostasis (GO: 0006884) | 0.404 | <0.01 |
| Vitamin K metabolic process (GO: 0042373) | 0.398 | <0.01 |
| Endosomal transport (GO: 0016197) | 0.392 | <0.01 |
| Histone exchange (GO: 0043486) | 0.387 | <0.01 |
| Primary metabolic process (GO: 0044238) | 0.382 | 0.01 |
| Barbed-end actin filament capping (GO: 0051016) | 0.348 | 0.02 |

**Supp. Info 1** Command line used for transcriptomic data analysis.

Quality check for every sequence in RNA-Seq datasets using program fastqc.

```
fastqc -t 6 RawSequence.fastq
```

Quality trimming and filtering using program cutadapt.

```
cutadapt -j 6 -m 50 -u 5 -U 5 -q 28,28 -o TrimmedSequence_F.fastq
-p TrimmedSequence_R.fastq RawSequence_F.fastq RawSequence_R.fastq
```

Build reference file for the mapping process using bowtie2-build.

```
bowtie2-build Reference.fasta ReferenceFile
```

Mapping of each sequence read to a reference file using program Tophat.

```
tophat -p 6 -
o ./tophat_output ./ReferenceFile ./TrimmedSequence_F.fastq ./Trim
medSequence_R.fastq
```

This code is to group similar unmapped sequences to examine the mapping process.

```
fastx_collapser unmapped.fasta | head -n 50
```

Sorting is required before the output bam file from the mapping process and convert into a sam file for further steps.

```
samtools sort -o accepted_hits_sorted.bam accepted_hits.bam

samtools view -o accepted_hits_sorted.sam accepted_hits_sorted.bam
```

Counting the total read abundance that mapped to the respective gene using program HTseq-count.

```
htseq-count -s no -t mRNA -i gene -m intersection-nonempty
accepted_hits_sorted.sam ReferenceFile.gff > ReadCount_sorted.csv
```
